# SLE Monocyte Subsets Are Pro-Inflammatory and Display Dysregulated Metabolism in Response to Bacterial Stimuli

**DOI:** 10.64898/2026.05.14.725094

**Authors:** Faye K Murphy, Anjali S Yennemadi, Natasha Jordan, Sarah Quidwai, Gina Leisching

## Abstract

Systemic lupus erythematosus (SLE) is associated with infection susceptibility and altered innate immune function. Monocyte metabolism is linked to appropriate cytokine release and bacterial containment. We investigated cytokine production and metabolic programming in the monocyte population from SLE patients and healthy controls following lipopolysaccharide (LPS) stimulation. SLE monocytes displayed increased IL-10, TNF, and IL-8 production, with impaired IL-1β induction. Metabolic profiling revealed altered substrate use, with increased glucose dependence and reduced fatty acid and amino acid oxidation after LPS stimulation. SLE patients exhibited reduced numbers of classical monocytes, expansion of intermediate monocytes, and dysregulated subset-specific metabolic reprogramming in response to LPS. This descriptive study provides a cornerstone for (i) understanding infection susceptibility in SLE, (ii) subset-resolved immunometabolic profiling as a tool in autoimmunity, and (iii) developing future metabolic-targeted therapeutic strategies

**Highlights:**

- Descriptive mapping shows SLE monocytes are proinflammatory with glucose dependence after LPS
- Classical and intermediate SLE subsets show divergent baseline metabolic preferences versus healthy
- SLE subsets display aberrant LPS responses, i.e.. increased glucose and reduced fatty acid oxidation
- This study provides a cornerstone for subset-resolved immunometabolism in infection susceptibility.

## Intro

Systemic lupus erythematosus (SLE) is a complex autoimmune disease driven by dysregulated immunity, aberrant immune cell metabolism and increased levels of circulating IFNα. Although therapeutic advances have improved life expectancy, SLE mortality rates remain threefold higher than in the general population, with infections as a leading cause of death[1, 2]. Despite optimised immunosuppressive therapies, SLE patients face heightened susceptibility to severe infections, spanning opportunistic pathogens, particularly bacteria[3]. However, whether this infection susceptibility reflects immunosuppression or an intrinsic defect in innate immune effector function remains unknown.

We now know that that the metabolic status of a cell can dictate its functional phenotype[4]. Metabolic alterations have emerged as an important feature of SLE monocytes. IFNα is a significant feature of SLE disease, and it has been implicated in impairing autophagy-mediated clearance of mtDNA, thereby promoting STING-dependent autoreactivity[5]. Accumulated mtDNA activates the cGAS-STING pathway, driving type I IFN responses alongside mitochondrial dysfunction, including reduced OXPHOS, decreased ATP, loss of membrane potential, and increased mROS, ultimately leading to mitochondrial damage and release of mitochondrial components[6, 7]. In fact, SLE monocytes have been associated with increased mitochondrial membrane potential, PINK1 mRNA, mtDNA content and JC1 aggregates[5]. Targeting metabolism can modulate this pathway, as mTOR inhibition with rapamycin reduces type I IFN production in SLE monocytes[8]. These observations raise the possibility that metabolic rewiring may underlie, or at least reinforce, the aberrant cytokine responses seen in lupus monocytes.

Monocytes are of particular interest because they rapidly migrate to sites of infection and phagocytose and destroy invading pathogens making them crucial in early host defence. In SLE, monocytes have been reported to exhibit an activated phenotype[9, 10] and secrete higher levels of pro-inflammatory cytokines[11, 12], suggesting that their ability to rapidly migrate to sites of infection may be compromised.

A further layer of complexity is provided by monocyte heterogeneity, with subsets differing in their migratory behaviour, inflammatory roles, and functional properties, and these subsets may be differentially involved in SLE pathogenesis. Classical monocytes, the most abundant subtype, have been shown to either differentiate into intermediate monocytes or undergo death by apoptosis or necrosis. Intermediate monocytes (CD14+CD16+) and non-classical monocytes (CD14loCD16+) are associated with patrolling behaviour, express the highest levels of antigen presentation related markers and possesses a greater ability to secrete inflammatory cytokines[13-15]. Understanding how these subsets differ metabolically is therefore important, particularly in a disease such as SLE where chronic inflammation may alter subset distribution and function. Research has shown that classical monocytes expressed higher levels of genes involved in carbohydrate metabolism, priming them for anaerobic energy production, nonclassical monocytes expressed higher levels of oxidative pathway components and showed a higher mitochondrial routine activity[16].

Despite growing interest in monocyte biology in SLE, relatively little is known about how metabolism is organised across ex-vivo monocyte subsets in health and disease. This is a noteworthy knowledge gap, because classical and intermediate monocytes likely rely on different metabolic programmes to support their distinct roles in immunity. Defining these subset-specific metabolic patterns may help explain how monocyte subsets contribute to the inflammatory phenotype of SLE and may identify new mechanisms linking immune activation to disease persistence. For these reasons, examining monocyte subset metabolism in SLE offers a useful route to understanding how cellular heterogeneity contributes to immune dysregulation in this disease. This descriptive study provides a cornerstone for (i) future hypothesis-driven mechanistic studies, (ii) the development of metabolic biomarkers for infection risk stratification, and (iii) the rational design of subset-targeted immunometabolic therapies in SLE.

## Methods

### Patient Recruitment

Peripheral blood was collected from 10 SLE patients and 10 healthy consented, age-matched individuals, and approved by the St James’s Hospital/Tallaght University Hospital Joint Research Ethics Committee (Table 1). Blood was collected in lithium-heparin–coated collection tubes and processed within the first hour. Full ethical approval for this study was granted by the St James’s Hospital/Tallaght University Hospital Joint Research Ethics Committee

**Table 1:** Donor demographics enrolled in this study. Values are n or median (interquartile range).

| Characteristics | SLE(N=10) | HC (N= 10) |
| --- | --- | --- |
| **<u>Demographics</u>** |  |  |
| *Age median, years (range)* | 36.5 (19-63) | 28.5 (24-38) |
| *Sex N* |  |  |
| Male | 1 | 3 |
| Female | 9 | 7 |
| *Race N* |  |  |
| Caucasian | 7 | 8 |
| Asian | 3 | 1 |
| African | 0 | 0 |
| Mixed | 0 | 1 |
| **<u>Lupus History</u>** |  |  |
| *Disease Severity N* |  |  |
| SLEDAI ≥ 4 | 4 | - |
| SLEDAI $<$ 4 | 6 | - |
| *Titres N* |  |  |
| Anti-Ro | 6 | - |
| Anti-Sm | 0 | - |
| Anti-dsDNA | 3 | - |
| ANA | 9 | - |
| RNP | 2 | - |
| *Medications N* |  |  |
| Mycophenolate mofetil | 4 | - |
| Mepacrine | 2 | - |
| Azathioprine | 1 | - |
| Hydroxychloroquine | 7 | - |
| Prednisolone | 2 | - |
| Warfarin | 1 | - |
| Rituximab | 1 | - |
| Belimumab | 1 | - |

### Cell Isolation and Stimulation

Peripheral blood mononuclear cells (PBMC) were isolated from whole blood by density centrifugation over Lymphoprep (STEMCELL Technologies). Cells were washed in PBS and resuspended in 3 mL serum-free RPMI (Gibco GlutaMAX) and layered onto 10 mL of hyperosmotic Percoll solution (GE Healthcare; 48.5% Percoll, 41.5% sterile H2 O, 0.16 M NaCl) to enrich for monocytes. Cells (0.5 × 10^6^ cells/mL) were plated on non-treated plates (Costar). The monocytes were then immediately stimulated with either 10ng/ml LPS or RPMI (‘unstimulated’) and placed in a humidified incubator at 37°C, 5% CO2 for 3 hours. Supernatants were harvested for cytokine quantification.

### Cytokine Measurement

The concentrations of IL-1β (BioLegend, 437016), IL-10 (BioLegend, 430604), IL-6 (BioLegend, 430516), IFN-γ (BioLegend, 430116), TNF (Invitrogen, 88-7346-88) were determined by ELISA following the manufacturer’s instructions.

### SCENITH (Single Cell mEtabolism by profiling Translation inHibition)

For flow cytometry analysis, 1 × 10^5^ cells per condition were washed with PBS and centrifuged at 400 x g for 5 min at room temperature (RT). The monocytes were then stimulated with either 10ng/ml LPS or RPMI (‘unstimulated’) for 3hrs at 37°C. Following stimulation, the monocytes were washed with PBS, centrifuged as mentioned earlier, and incubated with the following metabolic inhibitors for 40 min at 37°C according to the SCENITH protocol[17]: 2DG (100mM; ‘DG’), oligomycin (1μM; ‘Oligo’), a combination of 2DG and oligomycin (100mM, 1μM; ‘DGO’), or with harringtonine, a negative control (‘H’; 2ug/ml) or PBS as an untreated control (‘Co’). Puromycin (10μg/mL) was added for the final 35 minutes.

The monocytes were then surface stained for flow cytometry using fluorochrome-conjugated antibodies (Table 2) against CD14, CD15, CD16, CD3, CD56, CD19 (all 1:100), Fc block (1:50), and viability stain (1:200) in PBS for 20 min in the dark at RT. The cells were then washed, fixed and permeabilized using the Foxp3 staining kit (eBioscience #00-5523-00) as per manufacturer’s instructions. Cells were incubated for 30 mins at 4°C with anti-puromycin (1:100) and acquired on a Cytek Northern Lights (Cytek Biosciences). All analyses were performed using FlowJo v10.10 Software (BD Life Sciences).

**Table 2:**
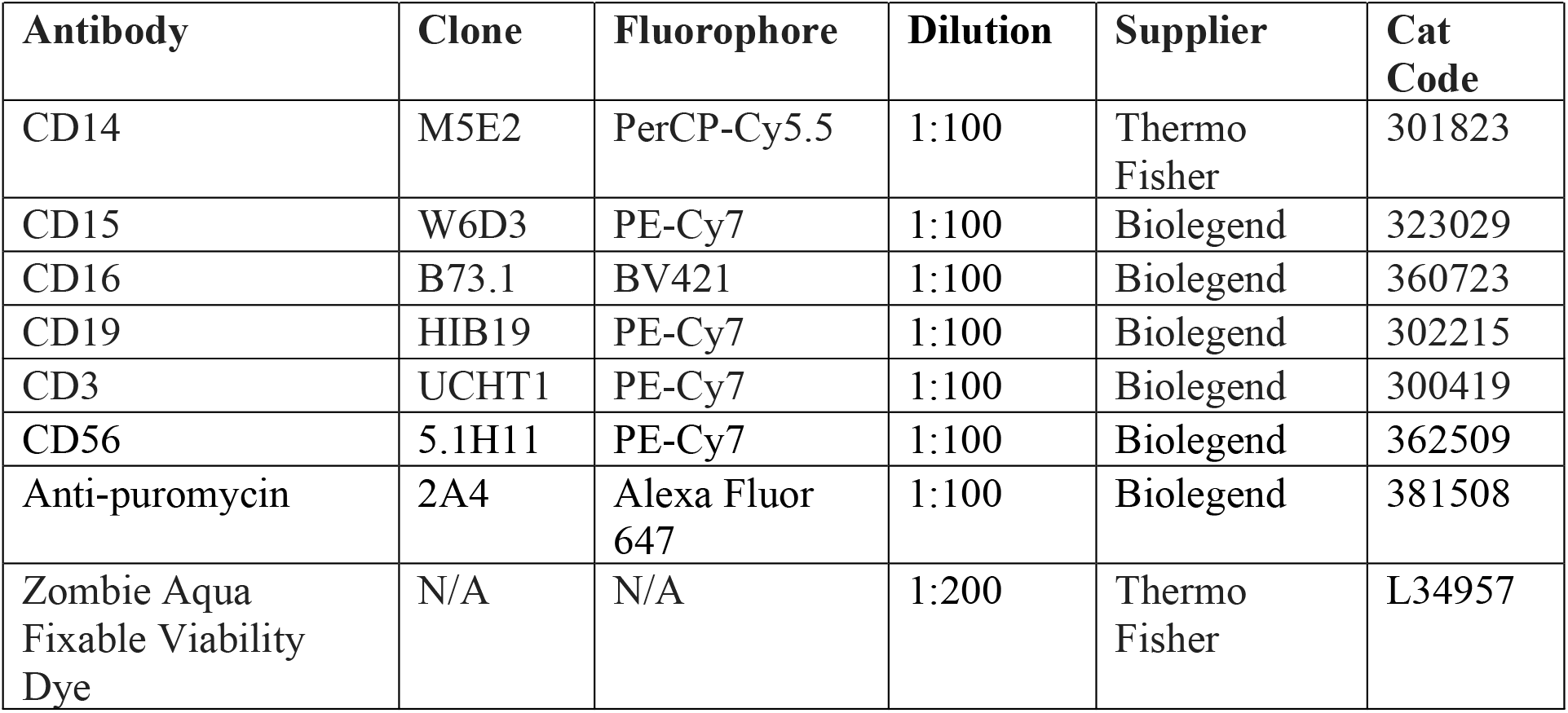
Antibodies, clones, and dilutions utilised in flow cytometry experiments.

The gating strategy involved removal of debris, doublets, and dead cells first, followed by gating on CD15, CD3, CD56 negative cells (Dump-) to eliminate potential cell contamination. Following this, monocytes were identified as CD14+. From the total monocyte population (CD14+), classical monocytes were defined as CD14+CD16-, intermediate monocytes as CD14+CD16+ and non-classical monocytes as CD14loCD16+.

### Statistical Analysis

Statistical analyses were performed using GraphPad Prism 10 software (GraphPad Software). Statistically significant differences between two unpaired non-parametric groups were determined using Mann Whitney tests with 2-tailed P values. Differences between 3 or more unpaired non-parametric groups were determined by RM 2-way ANOVA with uncorrected fisher’s LSD comparisons tests. P values of <0.05 were considered statistically significant.

## Results

### Dysregulated cytokine and metabolic responses in SLE monocytes after LPS stimulation

First, we wanted to functionally and metabolically characterise the total monocyte population. We observed no difference in the number of monocytes between healthy controls (HC) and SLE patients (Fig. 1A). Whether monocytes are increased or decreased in SLE is debated in the scientific community[18, 19]. While both HC and SLE monocytes enhance their IL-6 and IL-10 cytokine expression in response to LPS, the level of IL-10 secreted from LPS-stimulated SLE monocytes was significantly higher than healthy controls (Fig. 1B, 1C). On the other hand, HC monocytes can increase IL-1β in response to LPS, a response not observed in SLE monocytes (Fig. 1D). TNF and IL-8 are increased in SLE monocytes compared to healthy controls when stimulated with LPS, and at basal levels in IL-8 (Fig. 1E, 1F). These results suggest there is a dysregulated balance between pro- and anti-inflammatory cytokines in these SLE monocytes upon LPS challenge (Fig. 1G).

**Figure 1.**
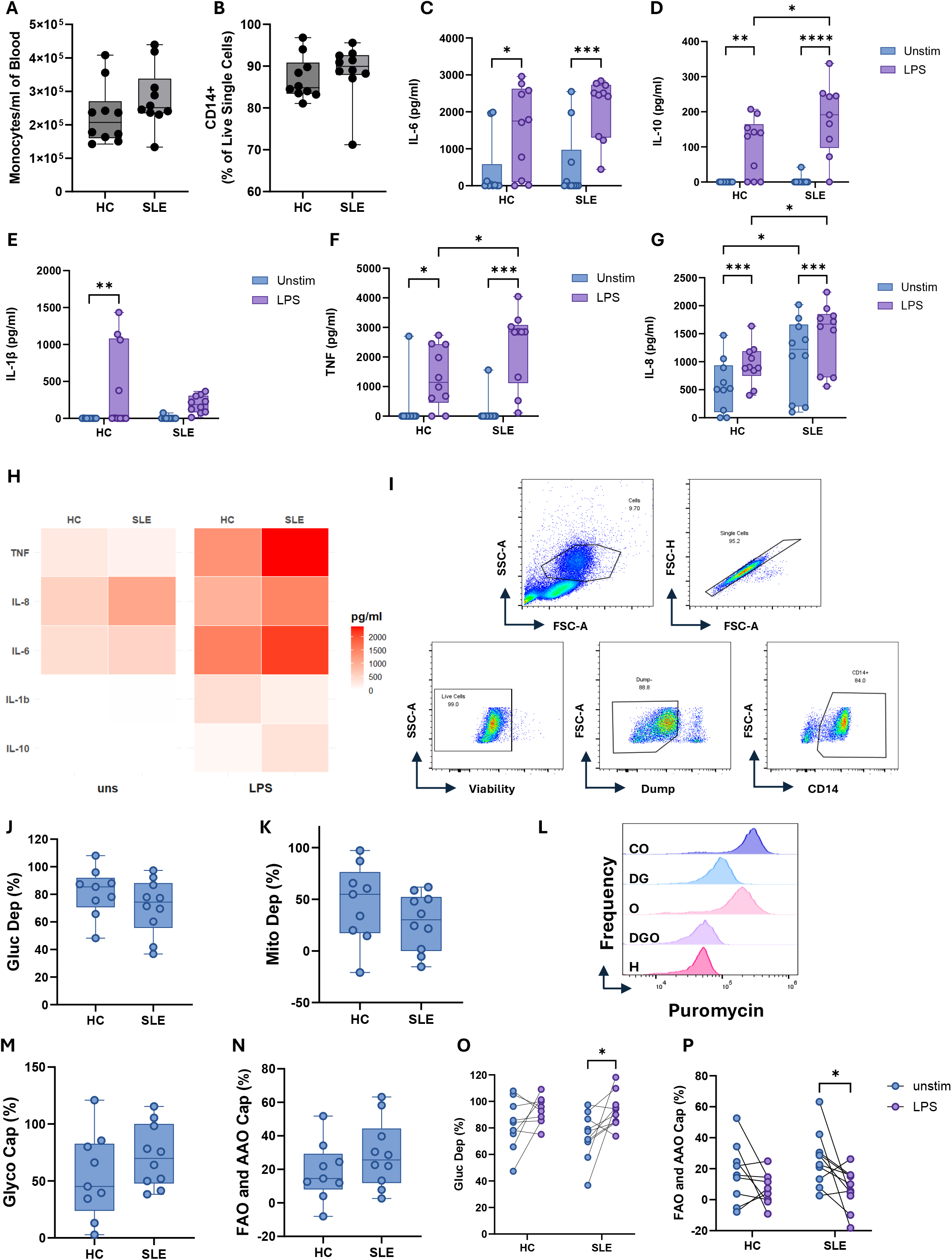
Dysregulated cytokine and metabolic responses are seen in SLE monocytes after LPS stimulation. Peripheral mononuclear cells were isolated from whole blood and stained with fluorochrome-conjugated antibodies against CD14, CD15, CD16, CD3, CD19, CD56 and analysed by flow cytometry or treated with metabolic inhibitors and puromycin for 40 min prior to staining. (A) Total monocyte counts per ml of whole blood. (B) Percentage of live single cells that are CD14+. The concentrations of IL-6 (C), IL-10 (D), IL-1β (E), TNF (F) and IL-8 (G) were assessed using ELISA and presented as pg/ml. (H) Heatmap showing the average pg/ml of each cytokine secreted from our cohort. (I) Representative flow cytometry gating strategy for identifying live, single, monocytes. Percent (J) glucose dependency, (K) mitochondrial dependence, (M) glycolytic capacity, and (N) fatty acid and amino acid oxidation of CD14+ monocytes (n = 8-10). (L) Representative histogram of puromycin MFI when CD14+ monocytes were treated with control or metabolic inhibitors. Percent (O) glucose dependency and (P) fatty acid and amino acid oxidation of CD14+ monocytes in response to LPS. P values calculated using Mann Whitney tests or two-way ANOVA with uncorrected Fisher’s LSD multiple comparison test.

With cytokine production being metabolically controlled[20], we investigated whether there are metabolic differences between HC and SLE monocytes using SCENITH[17]. In order to look at total monocytes, we defined monocytes as live, single, CD3-CD56-CD19-CD15- CD14+ cells (Fig. 1H). There was no difference in basal metabolism between HC and SLE monocytes (Fig. 1I, 1J, 1L, 1M). In response to LPS, SLE monocytes show increased glucose dependency compared to unstimulated SLE monocytes (Fig. 1N). This response was not seen in the healthy monocytes. While there is no difference in mitochondrial dependency or glycolytic capacity between healthy or SLE monocytes (Fig. S1E, Fig. S1F), fatty acid oxidation and amino acid oxidation in reduced in SLE monocytes when stimulated with LPS (Fig. 1O).

Together, these findings suggest that SLE monocytes display a distinct inflammatory and metabolic response to LPS, consistent with dysregulated immunometabolic programming. This suggests that SLE monocytes may be more metabolically flexible, allowing hyper- esponsiveness to bacterial infections, a mechanism often seen in SLE patients.

### Subset-specific metabolic reprogramming of monocytes in SLE

As mentioned earlier, monocyte subsets show altered phenotypes, with intermediate monocytes illustrating a mature phenotype that is associated with cell-to-cell adhesion, cell trafficking, proliferation, and differentiation[14, 15], we went on to investigate whether monocyte subsets show altered metabolism To separate total monocytes by flow, we defined classical monocytes as CD14+CD16-, intermediate monocytes as CD14+CD16+ and non-classical monocytes as CD14loCD16+ (Fig. 2A). We found that classical monocytes are reduced, and intermediate monocytes are expanded in SLE when compared to healthy controls (Fig. 2B). This agrees with the literature, also showing increased intermediate monocytes in SLE and other chronic diseases[21].

**Figure 2.**
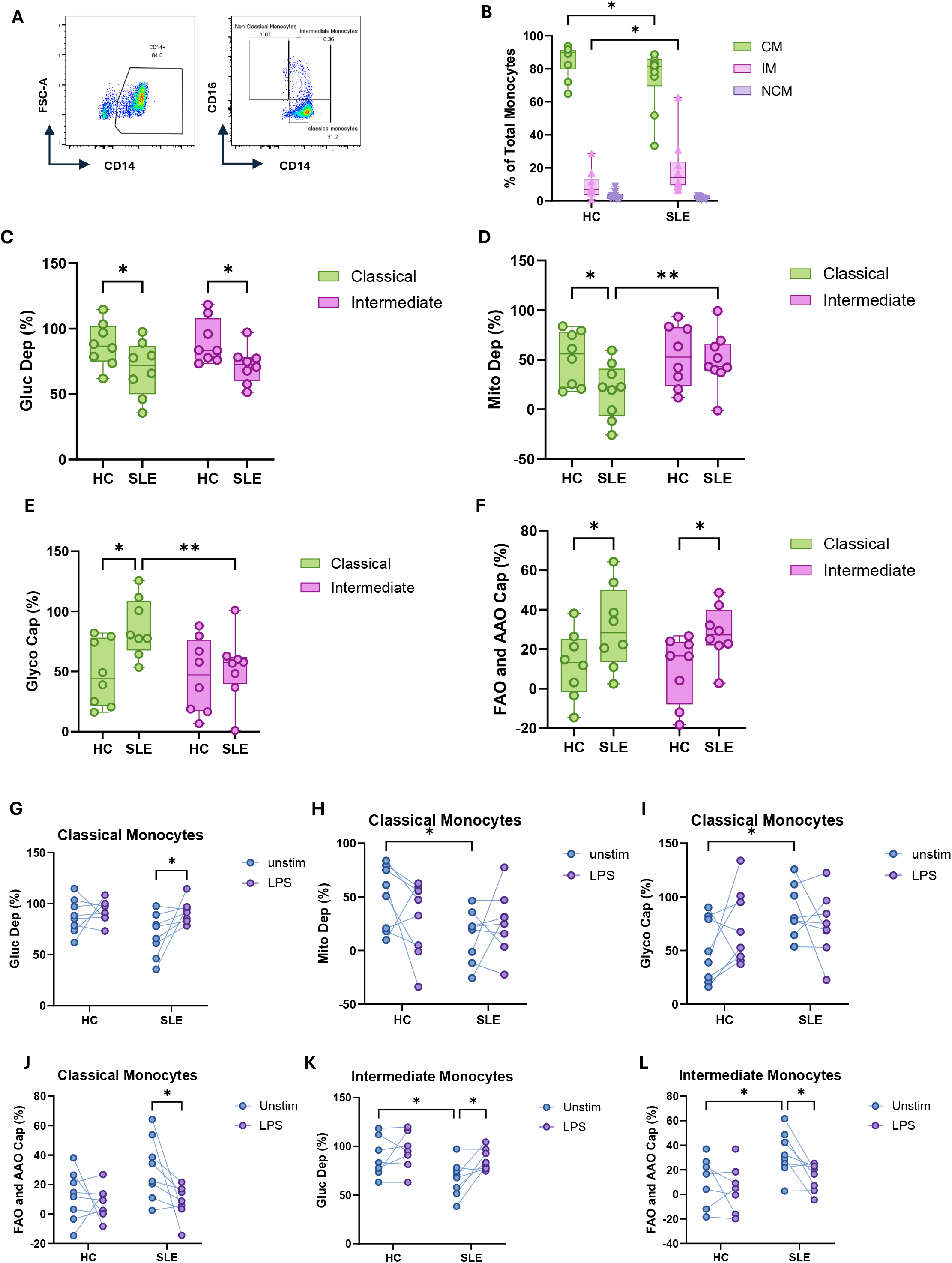
Subset-specific metabolic reprogramming of monocytes in SLE. Peripheral mononuclear cells were isolated from whole blood and stained with fluorochrome-conjugated antibodies against CD14, CD15, CD16, CD3, CD19, CD56 and analysed by flow cytometry or treated with metabolic inhibitors and puromycin for 40 min prior to staining. (A) Representative flow cytometry gating strategy for identifying classical and intermediate monocytes from CD14+ monocytes. Percentage of (B) classical, intermediate, and non-classical monocytes as a frequency of total live CD14+ monocytes. Percent (C) glucose dependency, (D) mitochondrial dependence, (E) glycolytic capacity, and (F) fatty acid and amino acid oxidation of classical and intermediate monocytes. Percent (G) glucose dependency, (H) mitochondrial dependence, IE) glycolytic capacity, and (J) fatty acid and amino acid oxidation of classical monocytes in response to LPS. Percent (K) glucose dependency and (L) fatty acid and amino acid oxidation capacity of intermediate monocytes in response to LPS (n = 8). P values calculated using two- way ANOVA and uncorrected fisher’s LSD multiple comparison test.

Due to low cell number of non-classical monocytes in SLE patients, they were excluded from our analysis. The frequency of each monocyte subset did not change with any metabolic inhibitor treatment (Fig. S1A, S1B), and baseline protein translation was equal in HC and SLE monocyte subsets (Fig. S1C, S1D). We observed that classical and intermediate monocytes from SLE patients show decreased glucose dependency compared to their HC counterparts (Fig. 2C). Additionally, intermediate monocytes from SLE patients show increased mitochondrial dependency compared to classical monocytes (Fig. 2D). SLE classical monocytes have reduced mitochondrial dependence compared to healthy classical monocytes (Fig. 2D). As a result, there is increased glycolytic capacity in SLE classical monocytes when compared to their healthy counterparts, and lower glycolytic capacity in SLE intermediate monocytes when compared to their classical monocyte counterparts (Fig. 2E). This may be due to intermediate monocytes being more mature and potentially less metabolically flexible than classical monocytes. This agrees with *Schmidl et al’*s finding that classical monocytes have higher levels of glycolytic genes while intermediate and non-classical monocytes have higher levels of genes associated with oxidative phosphorylation[16]. Additionally classical and intermediate monocytes from SLE patients have higher FAO and AAO capacity than healthy controls (Fig. 2F).

Next, we stimulated our monocytes with LPS to mimic bacterial infection to understand the immunometabolic changes of monocyte subsets in response to a bacterial infection. The glucose dependency of SLE classical monocytes significantly increases with LPS stimulation, and therefore reduced FAO and AAO capacity, which is not seen in HC monocytes (Fig. 2G, 2J). Although there is no significant change in response to LPS, the mitochondrial dependency of SLE classical monocytes trends upwards with stimulation, while the mitochondrial dependency of HC classical monocytes trends downwards with stimulation (Fig. 2H, Fig. 2I). This downward trend is seen in HC and SLE intermediate monocytes (Fig. S1G, S1H), suggesting SLE classical monocytes are uniquely altering their metabolism to response to bacterial stimuli. LPS stimulated SLE intermediate monocytes also increase their glucose dependency, and therefore, decrease their FAO and AAO capacity, which is not seen in LPS stimulated HC intermediate monocytes (Fig. 2K, 2L). As glycolysis is essential for the cytokine secretion from monocytes in response to LPS[4], this may be the reason why SLE monocytes have heightened pro-inflammatory cytokine secretion compared to HC in our cohort.

## Discussion

Our study addresses an important gap in the literature by defining how monocyte function and metabolism differ across subsets in SLE, rather than treating monocytes as a single population.

Our research is consistent with previous reports that SLE monocytes secrete higher levels of IL-6 and IL-10, though this response was evident only with LPS stimulation. IFNα has been implicated in driving IL-1β secretion[22], as a result, we would expect SLE monocytes to secrete higher concentrations of IL-1β, but this was not observed. Although all SLE monocytes were able to secrete IL-1β, some HC monocytes secreted higher levels than that seen from their SLE counterparts. In depth research into the TNF-TGF axis in SLE monocytes revealed that SLE monocytes secreted higher levels of TNF in response to apoptotic cells[23], we showed the opposite result in response to LPS. This implies that SLE monocytes are primed to induce a heightened TNF response to TLR7, TLR8 or TLR9, but not TLR4.

While research has shown higher levels of IL-8 in the serum of SLE patients and higher monocytic expression of CXCL8 compared to healthy controls, to our knowledge our finding of increased IL-8has not previously been shown[24, 25]. The heighted IL-8 secretion is biologically relevant as IL-8 is a neutrophil chemoattractant, that supports trapping bacteria and may contribute to the chronic inflammatory state in SLE. In this way, our findings fit with the broader literature on inflammatory monocyte dysregulation while revealing a more selective and stimulus-dependent cytokine profile.

A major strength and niche of our research is the integration of cytokine profiling with subset- specific metabolic analysis. Our work recapitulates the metabolic effects of LPS, specifically inducing metabolic reprogramming from oxidative phosphorylation to increased glycolytic capacity and glycolysis[26, 27]. This switch was seen in HC monocytes and SLE intermediate monocytes, but not in our SLE classical monocytes, with SLE classical monocytes showing increased glucose and mitochondrial dependency. This result may expose the hyper-activated state of classical monocytes often seen in SLE and help us understand why SLE monocytes can secrete higher levels of many cytokines. This altered metabolic phenotype in the subsets is essential to our understanding of monocyte biology and SLE pathogenesis. The lack of significance in our data may be due to insufficient monocyte adhesion and the associated metabolic programming that amplifies LPS responses[4].

This work and others show that intermediate monocytes are expanded in chronic inflammatory conditions[28, 29], but our results may help explain why this is the case. Their phenotype suggests a cell state that is poised for interaction with tissue environments, adhesion, and trafficking, but may also be metabolically specialised in a way that differs from classical monocytes. In SLE, the expansion of this subset, together with its altered response to LPS, may contribute to persistent low-grade inflammation and exaggerated responses to innate stimuli. The higher FAO and amino acid oxidation capacity observed in SLE classical and intermediate monocytes may also indicate compensatory use of alternative substrates, potentially reflecting adaptation to a chronically inflamed environment[30].

Overall, the present findings indicate that SLE monocytes display altered cytokine responses and subset-specific immunometabolic reprogramming. Rather than a simple increase in monocyte activation, SLE is characterised by a selective imbalance in pro- and anti- inflammatory cytokines, coupled with altered substrate preference and metabolic flexibility. This phenotype may be especially relevant in the context of bacterial infection, where LPS responsiveness could amplify inflammatory signalling and worsen disease flares. From a therapeutic perspective, these data support the possibility that targeting monocyte metabolism may help correct dysfunctional inflammatory responses in SLE.

Several limitations should be considered. First, our patient cohort was modest, and this study is intentionally descriptive, so these findings should be validated in a larger cohort and ideally stratified by disease activity, treatment exposure, and infection history. Secondly, SCENITH^™^ provides an informative snapshot of metabolic dependency, but it does not directly measure flux through specific pathways or mitochondrial respiration in real time. Additionally, subset analysis was restricted to classical and intermediate populations, and future work should examine non-classical monocytes as they also play important roles in both SLE and bacterial infections. Mechanistic studies are needed to determine whether the observed metabolic alterations are drivers of the cytokine phenotype or consequences of chronic inflammatory exposure. Finally, cytokine secretion is not the only role that monocytes play in bacterial infections, so understanding how these metabolic differences affect phagocytosis, antigen presentation and immunothrombosis is essential to further dissect monocytes in bacterial susceptibility.

In summary, our data support a model in which SLE monocytes are functionally reprogrammed towards an aberrant LPS response characterised by increased TNF and IL-8, altered IL-10 and IL-1β dynamics, and subset-specific changes in glucose use and alternative substrate oxidation. These findings strengthen the view that SLE is not only a disease of immune dysregulation, but also one of altered monocyte immunometabolism. This may be especially relevant during bacterial infection, when LPS exposure could amplify inflammatory signalling and contribute to flare risk.

---

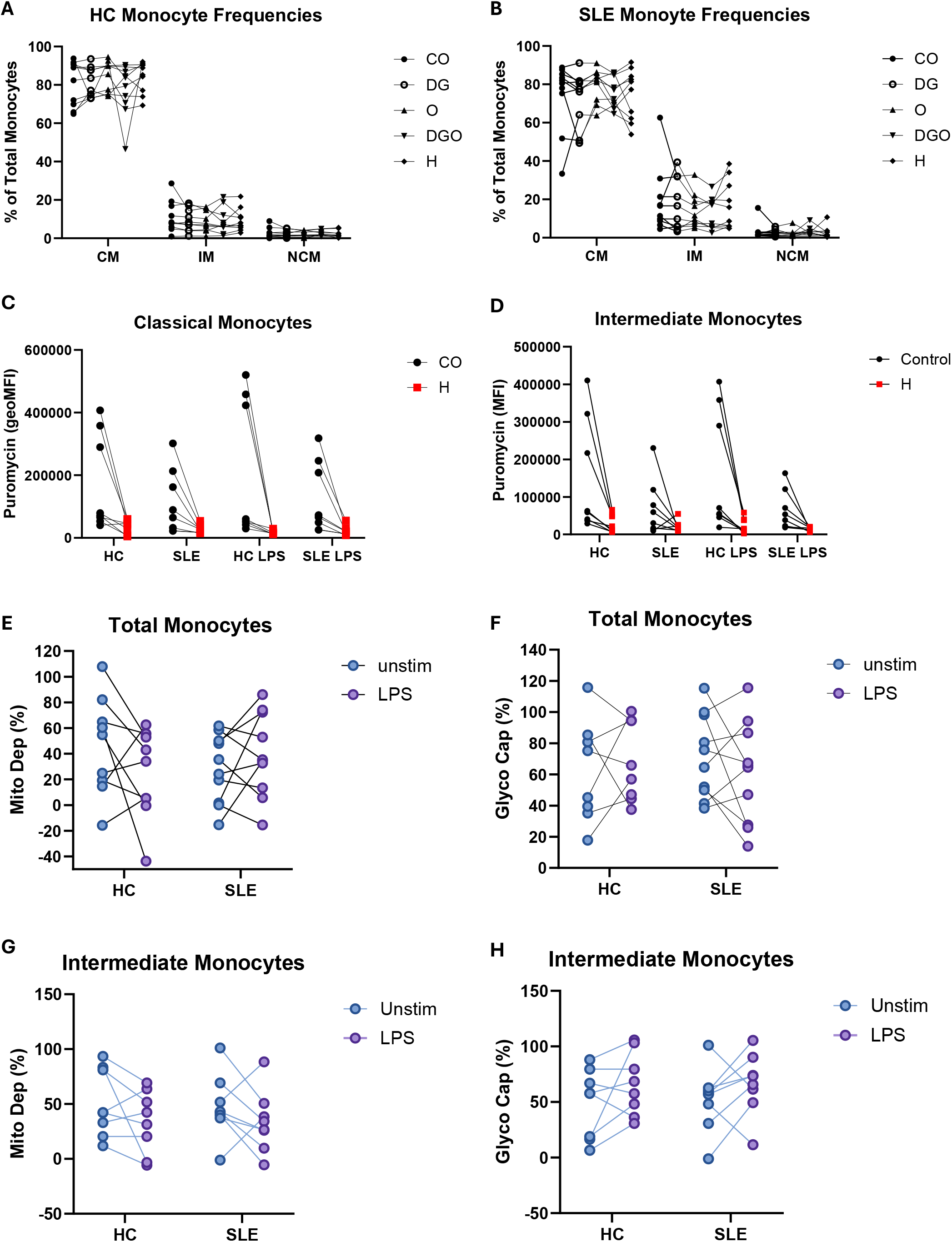

